# Two modes of PRC1-mediated mechanical resistance to kinesin-driven microtubule network disruption

**DOI:** 10.1101/2020.08.10.244491

**Authors:** April Alfieri, Ignas Gaska, Scott Forth

## Abstract

The proper structural organization of the microtubule-based spindle during cell division requires the collective activity of many different types of proteins. These include non-motor microtubule-associated proteins (MAPs) whose functions include crosslinking microtubules to regulate filament sliding rates and assembling microtubule arrays. One such protein is PRC1, an essential MAP that has been shown to preferentially crosslink overlapping antiparallel microtubules at the spindle midzone. PRC1 has been proposed to act as a molecular brake, but insight into the mechanism of how PRC1 molecules function cooperatively to resist motor-driven microtubule sliding and to allow for the formation of stable midzone overlaps has been lacking. Here we employ a modified microtubule gliding assay to rupture PRC1-mediated microtubule pairs using surface-bound kinesins. We discovered that PRC1 crosslinks always reduce bundled filament sliding velocities relative to single microtubule gliding rates, and do so via two distinct emergent modes of mechanical resistance to motor-driven sliding. We term these behaviors braking and coasting, where braking events exhibit substantially slowed microtubule sliding compared to coasting events. Strikingly, braking behavior requires the formation of two distinct high-density clusters of PRC1 molecules near microtubule tips. Our results suggest a cooperative mechanism for PRC1 accumulation when under mechanical load that leads to a unique state of enhanced resistance to filament sliding and provides insight into collective protein ensemble behavior in regulating the mechanics of spindle assembly.

## Introduction

Successful eukaryotic cell division requires the proper spatiotemporal organization of microtubule-based structures to segregate chromosomes and position the cell division plane (Kapoor, 2017). Specialized microtubule arrays, such as those assembled within the spindle midzone, undergo relative filament sliding at speeds that must be tightly regulated (McIntosh and Hays, 2016). Microtubule motions are primarily driven by motor proteins that crosslink and slide microtubules or exert pulling forces from filament ends (Forth and Kapoor, 2017). Filament motions are also modulated by non-motor microtubule-associated proteins (MAPs). MAPs can bundle microtubules into higher-order structural arrays, sort filaments by polarity, and regulate microtubule movement during sliding by producing brake-like resistance (Braun et al., 2011, 2017; Janson et al., 2007; Subramanian and Kapoor, 2012).

It is common for proteins that regulate microtubule motions and dynamics to function not as individual units but rather collectively as clusters of molecules, leading to behaviors that the constituent molecules do not exhibit alone. For example, the yeast kinesin-5 motor Cin8 walks towards microtubule minus-ends at the single particle level (Roostalu et al., 2011). However, when Cin8 forms clusters at microtubule minus-ends, these complexes can switch from fast minus-end to slow plus-end-directed motility and allow for the capture and sliding of antiparallel microtubules (Fallesen et al., 2017; Roostalu et al., 2011; Shapira et al., 2017). Another kinesin, the budding yeast kinesin-8 Kip3p, operates in a cooperative manner at microtubule plus tips to control filament depolymerization in a length-dependent manner (Varga et al., 2009, 2006). Dynamic microtubule plus-ends are decorated with EB1 ‘comets’ that saturate hundreds of binding sites at the growing tip, which likely enhances interactions with EB1 binding partners such as dynein (Bieling et al., 2008; Duellberg et al., 2014; Maurer et al., 2012; Pecreaux et al., 2006). In Physcomitrella patens, which like most land plants lacks cytoplasmic dyneins, none of the six minus-end-directed kinesin-14 members are processive as dimers (Jonsson et al., 2015). However, when artificially forced into ensembles, high processivity and fast motility are observed in the type-VI kinesin-14, providing the cells with a dynein-independent mechanism for retrograde transport. These observations suggest that localized high-density protein clusters can elicit functional mechanical outputs that are not generated by single molecules, but rather emerge via collective ensemble action.

Ensemble behavior is likely important for MAPs that crosslink microtubules within the mitotic spindle. One such MAP is PRC1, the human member of the conserved MAP65 family. Both PRC1 and the yeast MAP65 homolog Ase1 preferentially crosslink antiparallel microtubules both in vitro and in dividing cells (Bieling et al., 2010; Janson et al., 2007; Rincon et al., 2017; Subramanian et al., 2010; Tikhonenko et al., 2016; Yamashita et al., 2005). In metaphase, PRC1 has been shown to mediate the formation of ‘bridging fibers’ that crosslink sister kinetochore microtubule fibers (Kajtez et al., 2016; Polak et al., 2017; Vukušić et al., 2017). In anaphase, it has been suggested that PRC1 crosslinkers serve as mechanical linkages between segregating chromosomes and separating spindle halves. For example, in human cells PRC1 knockdown leads to reduced microtubule density at the spindle midzone, which results in the spindle halves being detached from each other (Mollinari et al., 2005, 2002; Zhu et al., 2006). Consistent with the model of PRC1 acting as a mechanical brake, increased chromosome and spindle pole segregation rates, as well as increased chromosome separation distances, are observed in PRC1-knockdown cells (Pamula et al., 2019). These data suggest that the collective actions of motor proteins external to the overlap pulling apart the microtubule bundle are resisted by PRC1 ensembles, though this behavior has not been specifically tested.

Several lines of evidence suggest that ensembles of MAP65 family members exhibit complex behaviors to resist microtubule sliding. For example, Ase1 has been shown to slow motor-driven microtubule sliding through an adaptive braking mechanism (Braun et al., 2011) which arises due to an entropic expansion force within densely crosslinked filaments that can prevent microtubules from sliding apart (Lansky et al., 2015). In vivo, Ase1 can selectively stabilize longer microtubule bundles while allowing for transport of shorter microtubules via motors, suggesting that Ase1 can produce overlap length-dependent resistive forces (Janson et al., 2007). Ase1 has also been shown to form multimers on single microtubules, where collisions between single diffusive Ase1 particles results in the formation of higher order complexes, though the functional role for this behavior is unclear (Kapitein et al., 2008). Similarly, direct mechanical measurements have revealed unique properties for the human crosslinker PRC1. Single molecule measurements of monomeric PRC1 constructs reveal the microtubule-binding domain alone can generate velocity-dependent frictional resistance when moved along the microtubule lattice (Forth et al., 2014). More recently, we have shown that ensembles of full-length PRC1 produce significant resistance against fast microtubule motions, but minimal resistance against slow motions (Gaska et al., 2020). Strikingly, no evidence of entropic expansion forces was detected for PRC1, suggesting a functional divergence from the yeast counterpart. Most intriguing is that when sliding is paused, PRC1 molecules undergo rearrangement to produce greater resistance upon resumption of sliding, suggesting the possibility that diffusion within overlaps results in the formation of higher-order PRC1 clusters, though such clusters were not directly observed (Gaska et al., 2020). Taken together, these findings suggest that PRC1 may form higher order structures with synergistic properties. However, it remains unclear how high density PRC1 clusters might form and how they might affect the mechanics of motor-driven filament pair separation.

To address these outstanding questions, we have employed a modified microtubule gliding assay using PRC1-crosslinked bundles rather than single microtubules. We imaged individual microtubule motions along with the distribution of PRC1 molecules in overlaps as bundles are ‘ruptured’ by the action of surface-bound kinesin motors which exert large forces and move microtubules with their minus-ends out. We find that PRC1 slows the separation of bundles under all conditions, but that a unique arrangement of PRC1 clusters near the filament tips occurs around half of the time and produces a substantially greater resistive force. We term the two observed modes of PRC1 resistance coasting and braking, and demonstrate that high density clusters allow for an increase in crosslinker persistence time in overlaps as well as a unique mechanical state that is capable of generating enormous resistive loads against filament separation.

## Results

### PRC1 Bundles Microtubules and Impedes Motor-Driven Sliding

To understand how microtubule bundles crosslinked by ensembles of PRC1 molecules resist sliding forces produced by motor proteins, we employed a modified microtubule sliding assay. First, a passivated coverslip sparsely decorated with antibodies to a 6-His motif was prepared in a sample chamber, and truncated kinesin-1 (K439) constructs with C-terminal 6-His tags were introduced and bound to the surface with their motor domains facing into solution. Next, 2-8 nM of a GFP-tagged PRC1 construct (hereafter referred to as PRC1) and microtubules were incubated together for ~20 minutes in 1X BRB80. These PRC1-mediated microtubule bundles were then introduced into the flow chamber along with either 10, 100, or 500 μM ATP. Over time, microtubule bundles landed on the kinesin-coated surface, at which point bundle disruption via motor-driven sliding began and could be visualized using TIRF microscopy (Figure 1A). Though we occasionally observed both single microtubules and bundles containing three or more microtubules, our analyses focus exclusively on pairs of microtubules as determined by fluorescence intensity from the microtubule imaging channel (Figure S1A). In each antiparallel pair of microtubules, our kinesin-1 construct walked toward the plus-ends, bringing them closer, and pushed the minus ends outward which resulted in a decrease of the overlap length. (Figure 1B). This assay allowed us to track microtubule movement and PRC1 distribution within microtubule overlaps over time.

**Figure 1:**
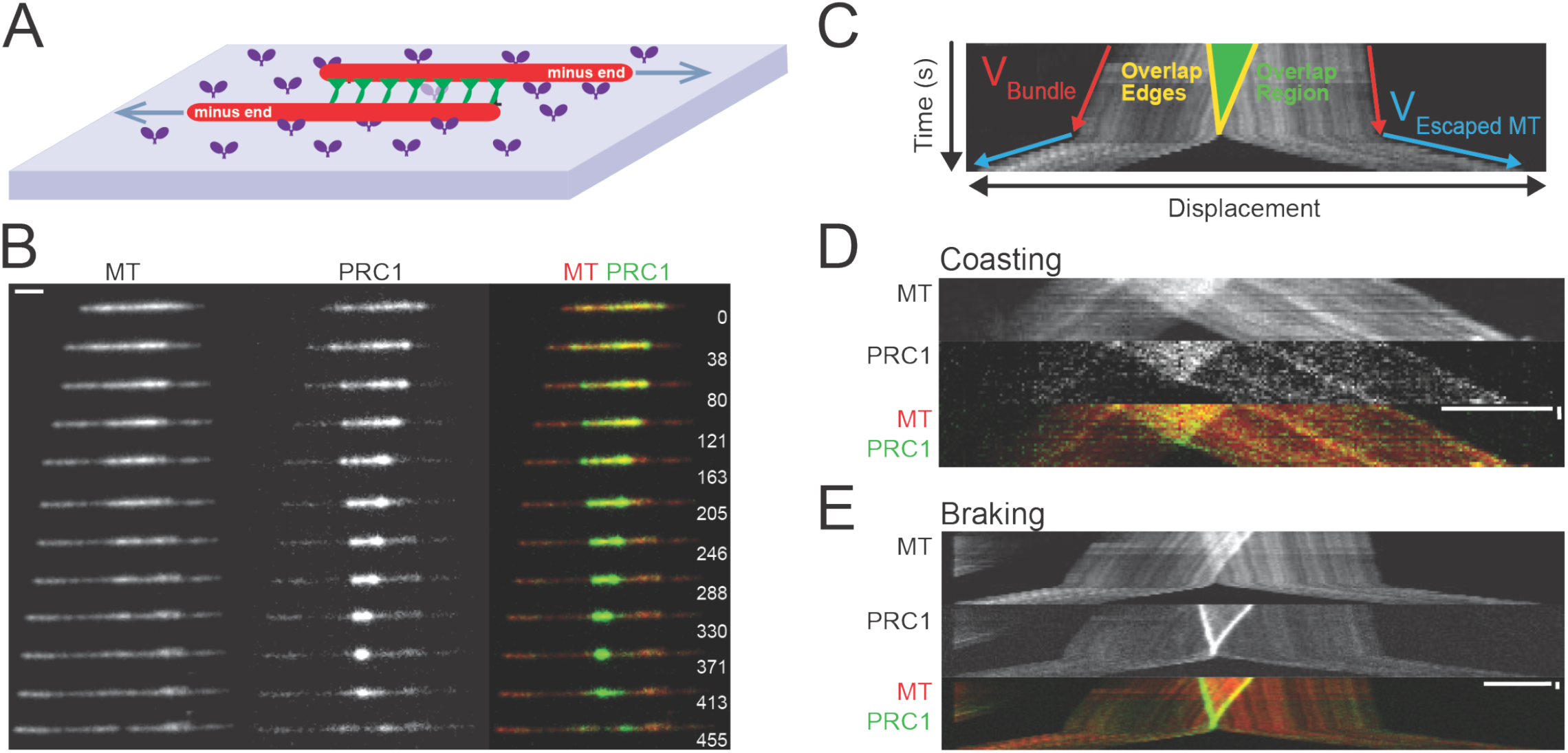
An Assay for Measuring Motor-driven Disruption of PRC1-crosslinked Microtubule Bundles. (A) Assay schematic. PRC1 (green)-mediated microtubule (red) bundles are disrupted by surface-bound, plus-end-directed kinesins (purple). (B) Time-lapse TIRF images of bundle disruption. Left: microtubules. Middle: PRC1. Right: Composite (microtubules in red, PRC1 in green). Frame acquisition time (in seconds) is shown on the right. (C) Example kymograph showing a PRC1-bundled microtubule pair sliding apart. Time (Y-axis) increases from top to bottom. The overlap region is colored green. Yellow lines depict overlap edges. Red arrows depict microtubule minus-ends used to calculate bundled velocities. Blue arrows depict microtubule minus-ends used to calculate escaped microtubule velocities. (D) Coasting kymograph. Top: microtubule. Middle: PRC1. Bottom: Composite (microtubules in red, PRC1 in green; X scale: 5 μm, Y scale: 20 sec). (E) Braking kymograph. Top: microtubule. Middle: PRC1. Bottom: Composite (microtubules in red, PRC1 in green; X scale: 5 μm, Y scale: 20 sec).

For each observed event we generated kymographs to visualize the time course of bundle disruption (Figure 1C). Each kymograph depicts one pair of microtubules sliding past each other in opposite directions with minus-ends leading. The microtubules pairs are crosslinked by PRC1 in the overlap region, which appears as a “V” shape in the middle of the kymograph. We calculated the instantaneous velocity from the slope of the microtubule edges throughout the course of bundle disruption and observed two distinct regions with different mean velocities: a slower velocity was measured while PRC1 was still crosslinking microtubules, thereby resisting sliding (“bundled”) and a faster velocity after the microtubules had separated from each other and PRC1 was no longer providing resistance via crosslinks (“escaped”, Figures S1B and S1C). For all measured events, we therefore parsed each microtubule’s motion into these two phases and calculated the average microtubule velocities for each. We determined that PRC1 was capable of slowing microtubule sliding across a range of kinesin stepping rates (Figure S1D). Surprisingly, we observed that some bundle disruption events exhibited little difference between bundled and escaped velocities (Figure 1D), while others had a much more significant difference (Figure 1E). Together, these data indicate that kinesin-driven microtubule gliding was impeded when microtubules were crosslinked by PRC1, but returned close to single filament sliding rates upon bundle disruption and the magnitude of velocity reduction appeared to vary between different bundles.

### PRC1 Resistance to Motor-Driven Microtubule Sliding Occurs in Two Distinct Phases

To quantify the observed differences in bundle behavior, we calculated the ratio of the bundled velocity (v_Bundle_) to the escaped microtubule velocity (v_Escaped_) for each microtubule in a bundle. This value provides a measure of how much microtubules are being slowed by PRC1 crosslinkers and was used as a criterion for categorizing events. Plotting all such ratios as a histogram revealed two distinct peaks. For approximately half (57%) of the events, bundled microtubule velocities were less than 40% of the escaped microtubule velocity, while for the remainder of the events bundled microtubule sliding velocities were > 40% of the escaped microtubule velocity (Figure 2A). We termed these behaviors braking and coasting, respectively. The ratio values and relative distribution of events were consistently observed across the range of ATP concentrations (Figure 2B).

**Figure 2:**
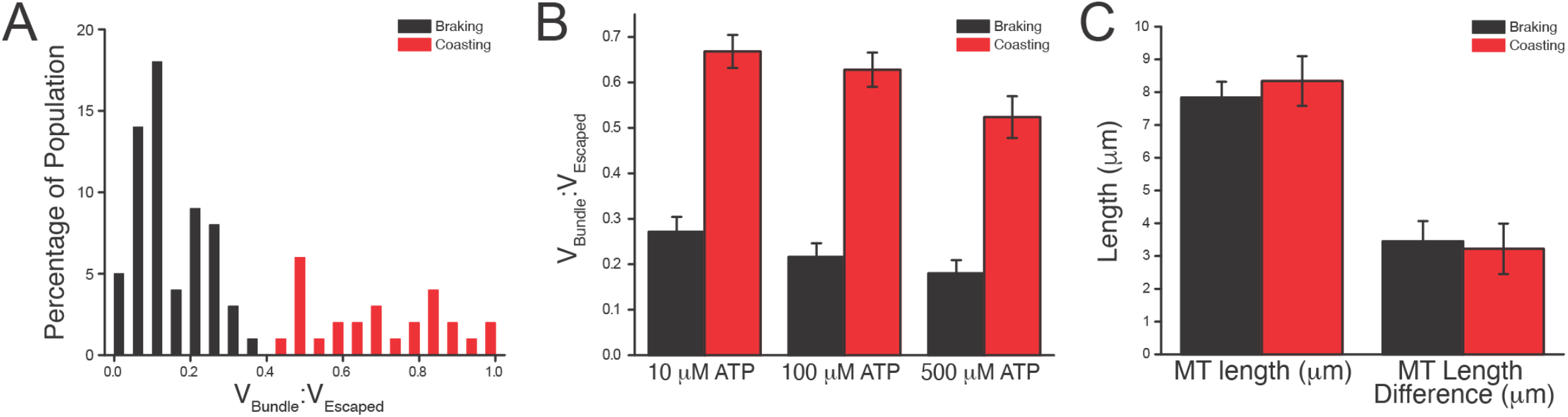
Two Emergent Modes of Mechanical Resistance to Bundle Disruption are Independent of Microtubule Geometry. (A) Distribution of the ratio of the bundled velocity to the escaped velocity (v_Bundle_:V_Escaped_) across all events at 100 μM ATP. Braking events (black, N = 62) occur at ratios <0.4 and coasting events (red, N = 27) occur >0.4. (B) Braking and coasting ratios calculated over a range of ATP concentrations. For braking events, N = 15 (10 μM ATP), 39 (100 μM ATP), 16 (500 μM ATP). For coasting events, N = 16 (10 μM ATP), 24 (100 μM ATP), 11 (500 μM ATP). (C) Comparison of the average microtubule lengths in bundles and the average difference in microtubule lengths within a bundle for braking (black) and coasting (red) events. N = 57 (braking microtubule lengths), 36 (coasting microtubule lengths), 29 (braking microtubule length difference), 18 (coasting microtubule length difference). All error bars are SEM.

To better characterize these two modes of behavior, we looked for potential differences in the construction of the bundles. First, we examined microtubule length, conjecturing that bundles containing longer microtubules might favor one state over the other, but found no significant difference between microtubule lengths within bundles from braking or coasting events. Additionally, when the relative difference in microtubule lengths within each bundled pair was calculated, we observed no difference between braking and coasting events (Figure 2C). We next compared bundled and escaped velocities for each disruption event and found that longer microtubules tended to have higher bundled velocities, but observed no microtubule-length dependence for the escaped velocities (Figure S2). We speculate that longer microtubules interact with a larger number of kinesin motors which leads to increased force production, therefore permitting the bundled microtubules to move at higher velocities even when impeded by PRC1 crosslinks. Based on these observations, we concluded that the microtubules themselves were not largely responsible for the differences between braking and coasting behaviors, and that these differences were instead likely an intrinsic property of PRC1.

### Braking Events Retain More PRC1 as the Overlap Shrinks than Coasting Events

To further probe the mechanistic differences between these two modes of microtubule behavior, we next analyzed the total amount of PRC1 in overlaps. We hypothesized that braking overlaps might simply contain more PRC1 than coasting overlaps, which would then impart a greater ability to resist microtubule sliding. To test this, we integrated the GFP-PRC1 intensity within the overlap region at each time point (Figure 3A). First, we compared the integrated GFP-PRC1 intensity in overlaps at T = 0, when bundles first make contact with the motors on the coverslip surface. While braking overlaps have a slightly higher average initial integrated intensity than coasting events (1432 au and 805 au, respectively), the majority of braking and coasting events fall into a similar range of initial integrated intensities (Figure 3B). Therefore, we next looked at the time-dependent change in the amount of GFP-PRC1 in the overlap region as microtubules slid apart. For populations of braking and coasting events with similar initial amounts of PRC1 in the overlap region (~900-1600 au), braking events retain more PRC1 molecules as the overlap shrinks. We observed a linear decrease in GFP integrated signal as overlap lengths shortened, consistent with loss of PRC1 crosslinkers from overlaps into solution. However, upon reduction to just 10% of the initial overlap length, braking events still retained ~50% of the molecules they started with, while coasting events only retained ~30% (Figure 3C). Therefore, we conclude that it is not the initial amount of PRC1 in overlaps but the retention of molecules that can distinguish braking from coasting behavior.

**Figure 3:**
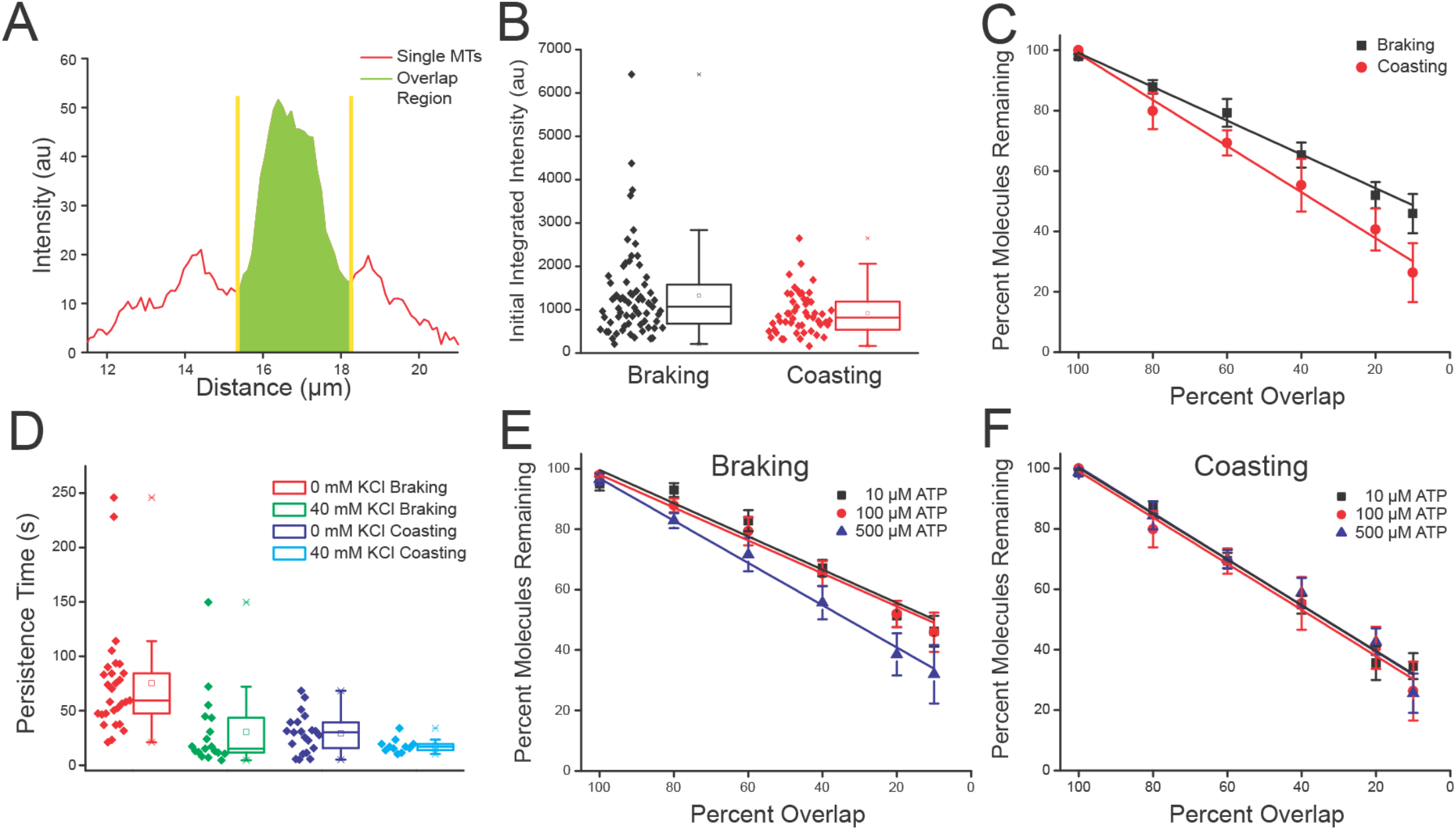
Retention of PRC1 Molecules Differs Between Braking and Coasting Events. (A) Sample GFP-PRC1 fluorescence intensity line scan of a bundle. The overlap region is highlighted in green, bounded by yellow lines. (B) Individual values (solid diamonds) and box plots of initial PRC1 integrated intensities measured from braking (black) and coasting (red) events. N = 69 (braking), 52 (coasting). (C) Percentage of PRC1 molecules remaining within the overlap region as function of overlap length for braking (black) and coasting (red) events at 100 μM ATP. Number of bundles averaged for each point (braking): 10 (100%), 11 (80%), 11 (60%), 11 (40%), 11 (20%), 8 (10%). Number of bundles averaged for each point (coasting): 8 (100%), 4 (80%), 6 (60%), 4 (40%), 4 (20%), 3 (10%). (D) Individual values (solid diamonds) and box plots of persistence times for braking (red: 0mM KCl, green: 40mM KCl) and coasting (purple: 0mM KCl, cyan: 40mM KCl) bundles at 100 μM ATP. N = 29 (braking, 0 mM KCl), 18 (coasting, 0 mM KCl), 21 (braking, 40 mM KCl), 11 (coasting, 40 mM KCl). (E) Percentage of PRC1 molecules remaining within the overlap region as function of overlap length for braking events at 10 (black), 100 (red), and 500 (blue) μM ATP. Number of bundles averaged for each point: 8, 10, 8 (100%), 6, 11, 7 (80%), 6, 11, 7 (60%), 8, 11, 7 (40%), 8, 11, 7 (20%), 5, 8, 4 (10%) for 10, 100, and 500 μM ATP, respectively. 100 μM data replotted from (C). (F) Percentage of PRC1 molecules remaining within the overlap region as function of overlap length for coasting events at 10 (black), 100 (red), and 500 (blue) μM ATP. Number of bundles averaged for each point: 8, 8, 5 (100%), 7, 4, 5 (80%), 7, 6, 4 (60%), 8, 4, 5 (40%), 6, 4, 4 (20%), 5, 3, 3 (10%) for 10, 100, and 500 μM ATP, respectively. 100 μM data replotted from (C). All error bars are SEM. Box plots depict mean, median, standard deviation, 95% confidence interval, and maximum/minimum values for each data set.

Expanding on these observations, we next hypothesized that decreasing PRC1’s lifetime on microtubules would potentially alter the mode of bundle disruption. PRC1’s interaction with and lifetime on the microtubule lattice can be modulated by changing the ionic strength of the buffer. Therefore, to test our hypothesis, we allowed PRC1 to bundle microtubules in 1X BRB80 buffer as before, but increased the KCl concentration in our final assay buffer to 40 mM. The duration of braking events at 40 mM KCl was lower than those without additional KCl. Coasting events at 40 mM KCl were still shorter-lived than braking events under the same conditions (Fig 3D). Increasing the ionic strength therefore not only decreases PRC1’s lifetime in bundles, but the lifetime of the bundles themselves. Taken together, these results suggest that increased PRC1 lifetime within overlaps was correlated with a preference for braking over coasting modes.

Next we wanted to see if these observations held true across different baseline microtubule sliding velocities by varying the ATP concentration available for the kinesin motors. PRC1 molecules in braking events at 100 μM ATP behave similarly to those at 10 μM ATP, while at 500 μM ATP PRC1 is more rapidly lost from the overlap region (Figure 3E). At higher microtubule sliding velocities, braking events behave more like coasting events, which lose PRC1 from the overlap region at a similar rate across all three ATP concentrations (Figure 3F). Additionally, the amount of PRC1 molecules remaining was similar for braking events at 0 and 40 mM KCl, but coasting events lost PRC1 molecules more rapidly when there was no additional KCl in the final buffer (Figure S3). Taken together, the retention of PRC1 molecules seems to be a contributing factor in bundle behavior, differing between braking and coasting conditions and dependent on ionic strength of the buffer.

### PRC1 Distribution Differs Between Braking and Coasting Events

We next hypothesized that the distribution of PRC1 in the overlap may be important to how certain bundles retain more PRC1 over time. Therefore, we compared the average variance of the GFP signal in the overlap region, where lower variance (Figure 4A) suggests more even distribution of PRC1 and a higher variance indicates regions of locally high and low PRC1 density within the overlap (Figure 4B). At 100 μM ATP, braking events have consistently greater variance in the GFP signal in the overlap region than coasting events. In braking events, the variance continuously increases as the overlap shrinks, while in coasting events the variance is lower and remains relatively constant up until bundle disruption (Figure 4C). Braking events across all three ATP concentrations show a similar pattern as in Figure 3B; 10 and 100 μM ATP braking events have greater overlap GFP variance than coasting events and both increase in variance, with 100 μM ATP braking events increasing more sharply at <20% of the initial overlap length. At 500 μM ATP, variance in braking events is again more similar to that of coasting events (Figure 4D). Variance in coasting event overlaps at all three ATP concentrations is similar up until 20% of the initial overlap length, but coasting events at each ATP concentration still have much lower variance in GFP intensity than braking events at 10 and 100 μM ATP (Figure 4E). Together these results suggest that the distribution of PRC1 plays a role in whether a bundle displays braking or coasting behavior.

**Figure 4:**
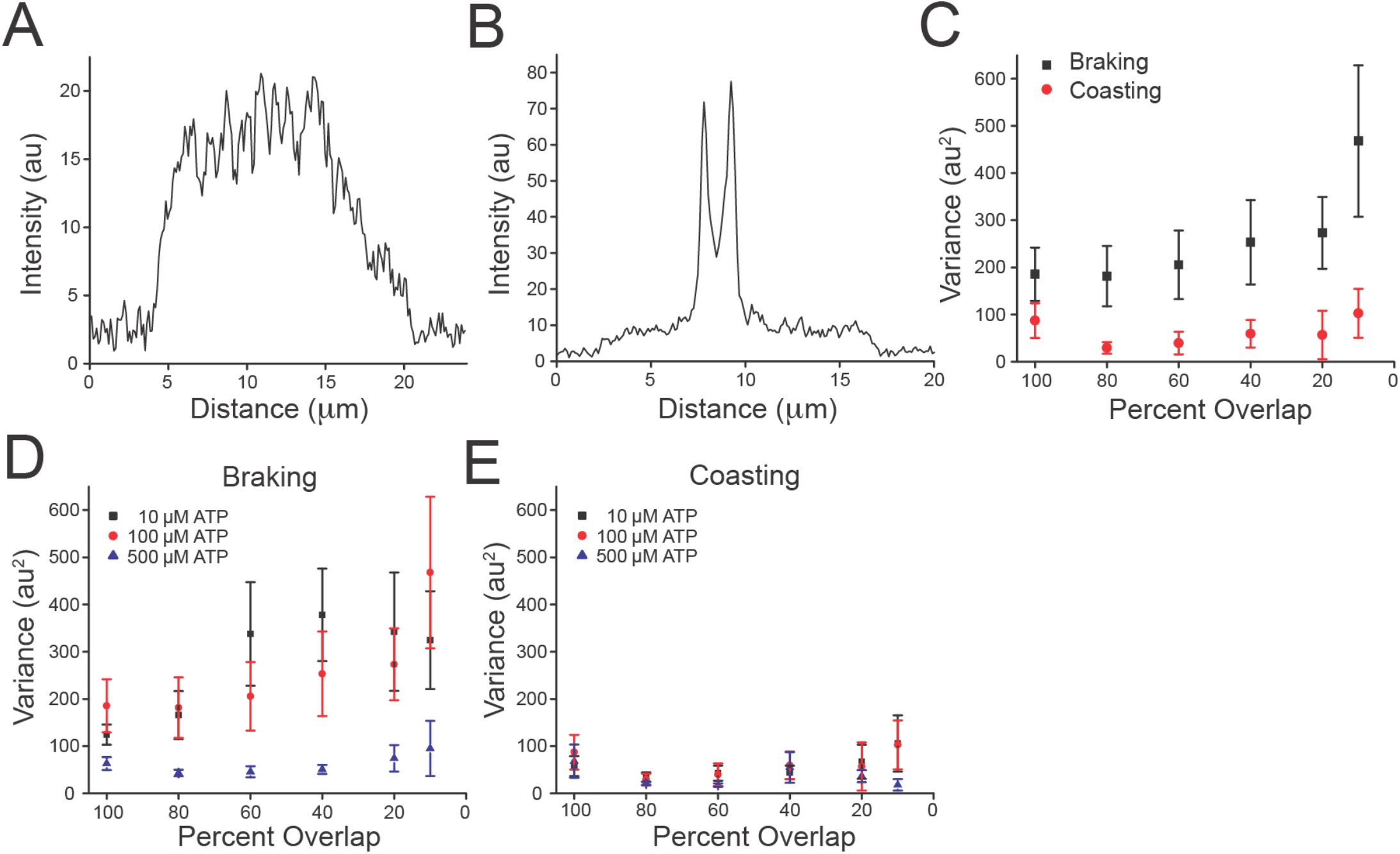
Distribution of PRC1 Differs in Braking and Coasting Events. (A) An example of PRC1 intensity within overlap region with low calculated variance. (B) An example of PRC1 intensity within overlap region with high calculated variance. Calculated initial integrated intensities for examples in (A) and (B) were within ~200 au of each other. (C-E) GFP-PRC1 fluorescence intensity variance in the overlap region as a function of decreasing overlap length. (C) Comparison of the GFP-PRC1 variance in braking (black) and coasting (red) events at 100 μM ATP. Number of bundles averaged for each point are the same as in 3C. (D) Comparison of the GFP-PRC1 variance in braking event overlaps at 10 (black), 100 (red), and 500 (blue) μM ATP. Number of bundles averaged for each point are the same as in 3E. 100 μM data replotted from (4C). (E) Comparison of the GFP-PRC1 variance in coasting event overlaps at 10 (black), 100 (red), and 500 (blue) μM ATP. Number of bundles averaged for each point are the same as in 3F. 100 μM data replotted from (D). All error bars are SEM.

### Braking Events Contain Two High-Density PRC1 Clusters at the Overlap Edges

The measured differences in variance in GFP-PRC1 signal in the overlap led us to carefully examine kymographs and time-lapse fluorescence intensity linescans for patterns in the variations in PRC1 distribution. We observed that braking events frequently contained two peaks of GFP-PRC1 signal at each of the overlap edges in kymographs that can be clearly by linescan analysis (Figure 5A-D, 1C). Coasting events lacked these two distinct peaks in fluorescence (Figure 5E-H). To confirm that the formation of these peaks was a result of motor-driven movement of microtubules, we replicated the assay but did not include ATP, which resulted in bundles bound to the surface whose microtubules remained static. We observed PRC1 distribution within bundles at two time points: immediately after flowing them into the chamber and after two hours. We then quantified the number of peaks present at these time points and found that the majority of bundles (~65%) contained no peaks initially, and this number increased to ~77% after two hours. About 23% of bundles initially contained one peak, and after two hours this decreased to 16%. Finally, ~13% of bundles contained two peaks initially, and this decreased to ~9% after two hours (Fig S5A). These data led us to conclude that in the majority of cases, the doublet peaks observed in braking events emerged due to motor-driven filament sliding and not time-dependent aggregation of PRC1.

**Figure 5:**
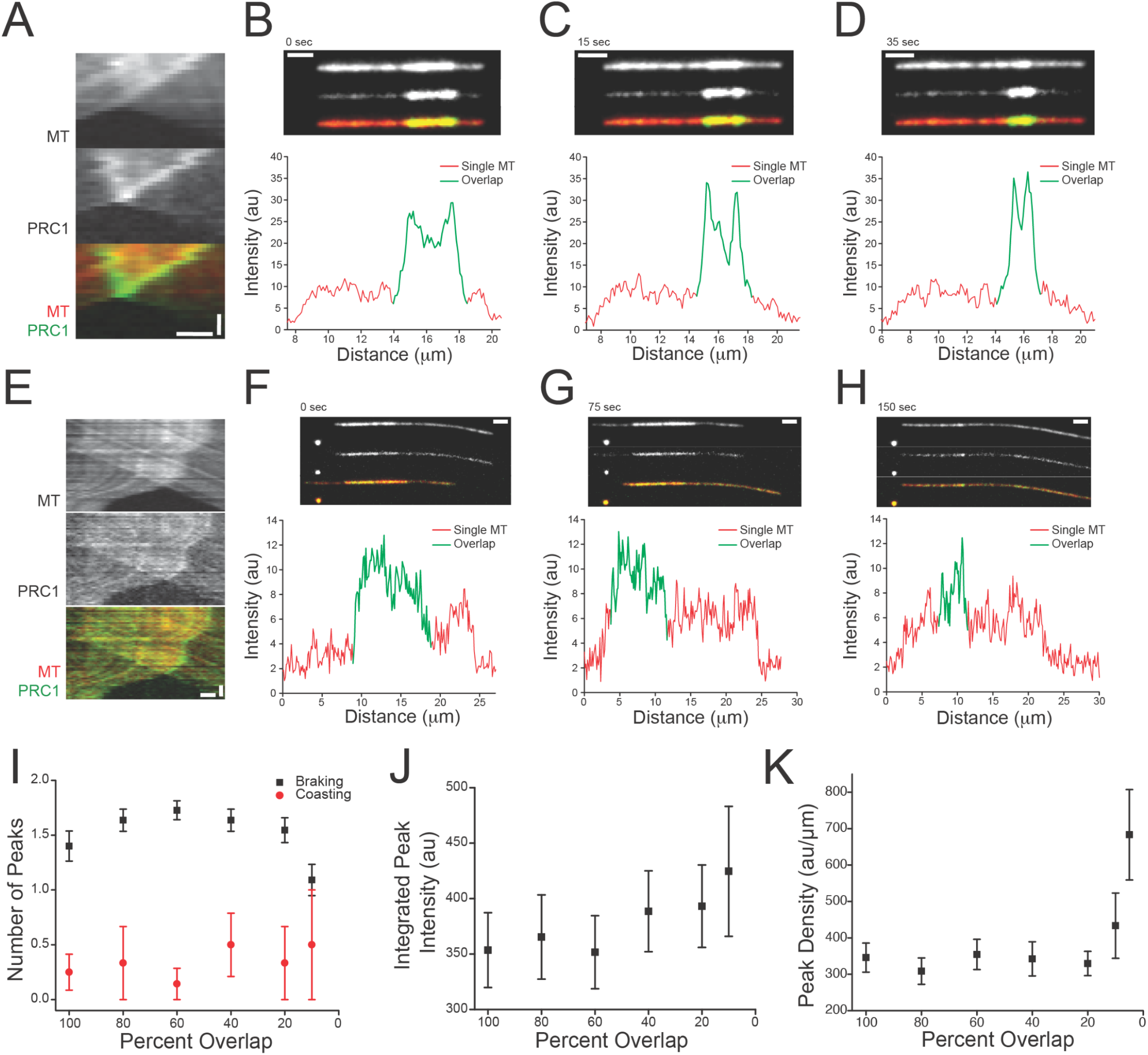
PRC1 Forms Two Tip-localized Clusters in Braking Event Overlaps That Accumulate Molecules and Increase in Density. (A-D) Example braking event kymograph and fluorescence intensity line scans showing PRC1 clusters/peaks in GFP-PRC1. (A) Overlap region of a braking event bundle. Top: microtubule. Middle: PRC1. Bottom: Composite (microtubules in red, PRC1 in green. X scale: 1 μm. Y scale: 25 sec). (B-D) Time-lapse fluorescence intensity line scans depicting the development of peaks in GFP intensity that correlate to overlap-edge clusters seen in (A). Single microtubule GFP signal in red, overlap GFP in green. Scale: 2 μm. (E-H) Example coasting event kymograph and fluorescence intensity line scans showing lack of clusters or peaks in GFP-PRC1. (E) Overlap region of a coasting event bundle. Top: microtubule. Middle: PRC1. Bottom: Composite (microtubules in red, PRC1 in green. X scale: 1 μm. Y scale: 25 sec). (F-H) Time-lapse fluorescence intensity line scans depicting more uniform GFP signal in the overlap region. Single microtubule GFP signal in red, overlap GFP in green. Scale: 2 μm. (I) Number of GFP-PRC1 peaks in braking (black) and coasting (red) events as function of decreasing overlap length at 100 μM ATP. Braking events have close to two peaks on average until distinct peaks can no longer be resolved at short overlap lengths (<20% of the initial overlap length). Coasting events have 0.5 peaks on average. Number of individual peaks included for each mean value (braking): 14 (100%), 18 (80%), 19 (60%), 18 (40%), 17 (20%), 12 (10%). Number of individual peaks averaged for each point (coasting): 8 (100%), 4 (80%), 6 (60%), 4 (40%), 4 (20%), 3 (10%). (J) Integrated intensity of GFP signal within peaks during braking events at 100 μM ATP. Number of individual peaks averaged for each point: (100%), 18 (80%), 19 (60%), 18 (40%), 17 (20%), 12 (10%), (5%). (K) GFP density (integrated intensity per micron) within braking event peaks at 100 μM ATP. Number of individual peaks averaged for each point: 10 (100%), 16 (80%), 14 (60%), 15 (40%), 17 (20%), 12 (10%), 9 (5%). All error bars are SEM.

Next, we quantified the time evolution of GFP fluorescence peaks in overlaps. At 100 μM ATP, the number of peaks within braking events increase from ~1.4 to 1.8 on average throughout the disruption event before rapidly decreasing to ~1.2 peaks as the clusters of PRC1 molecules merge just before disruption. In contrast, the number of observed peaks within coasting events increase from ~0.2 to 0.5 peaks on average (Figure 5I). Across ATP concentrations, braking bundles at 500 μM ATP take slightly longer to develop a second peak than those at 10 or 100 μM ATP, but all three conditions are similar after 80% of the initial overlap length (Figure S5B). For coasting events, the number of peaks varies between ~0.2 and 1 peak(s), with events at 10 and 500 μM ATP being the most similar, and events 100 μM ATP having fewer peaks on average (Figure S5C). From these results we conclude that the formation of multiple peaks near microtubule tips is essential to braking behavior.

Next we wanted to examine how these PRC1 clusters in braking events change over time, so we integrated the region surrounding the peak maxima. At 10 μM ATP, integrated braking event peak intensities increase from 300 to ~375 au and then remains fairly constant after 40% of the initial overlap length. At 100 μM ATP, peaks continuously increase from 300 to 475 au, and at 500 μM ATP, peak intensities fluctuate between 200 and 275 au (Figure 5J, S5D). We also calculated the PRC1 density in peaks using the full width at half-maximum (FWHM). At 100 μM ATP, peak densities remained fairly constant (300-400 au/μm) until <20% of the initial overlap length, where they increased rapidly to ~700 au/μm (Figure 5K). Increasing ionic strength in the final buffer resulted in braking events containing closer to one peak on average, and these singular peaks had lower integrated intensities (Fig S5E-F). Peak densities were lower than in the 0 mM additional KCl condition and remained consistent through the entire life of the overlap (~150-200 au/μm, Figure S5G). For coasting events, the number of peaks did not differ between buffer conditions (Fig S5H). Comparison of braking event peak heights at 50% of the initial overlap length for all ATP concentrations revealed a negative correlation between peak height and average bundled velocity. Higher bundled velocities, usually seen at 500 μM ATP, tend to have lower peak intensities, while the microtubule pairs that have lower bundled velocities have more intense peaks (Fig S5I). Taken together, these results suggest that the amount and distribution of PRC1 within peaks is an important determinant of braking behavior. Braking event peaks not only gain PRC1 molecules over time, but PRC1 density greatly increased, particularly at short overlap lengths.

## Discussion

We have employed in vitro reconstitution assays and TIRF microscopy time-lapse imaging to show that PRC1 exhibits two distinct behaviors when crosslinking two microtubules that are being slid apart by surface-bound kinesin molecules. We term these distinct states braking and coasting. In braking events, PRC1 slows microtubule movement significantly more than in coasting events relative to the single microtubule velocity. Braking events retain more molecules of PRC1 within overlaps during microtubule sliding than coasting events. Strikingly, bundles undergoing braking develop two dense clusters of PRC1 at the overlap edges that accumulate molecules as the overlap shrinks, whereas bundles undergoing coasting maintain a more uniformly distributed population of PRC1 (Figure 6). These data suggest that both the retention and relative distribution of PRC1 molecules within overlaps are critical to determining the mechanical function of sliding crosslinked filament pairs.

**Figure 6:**
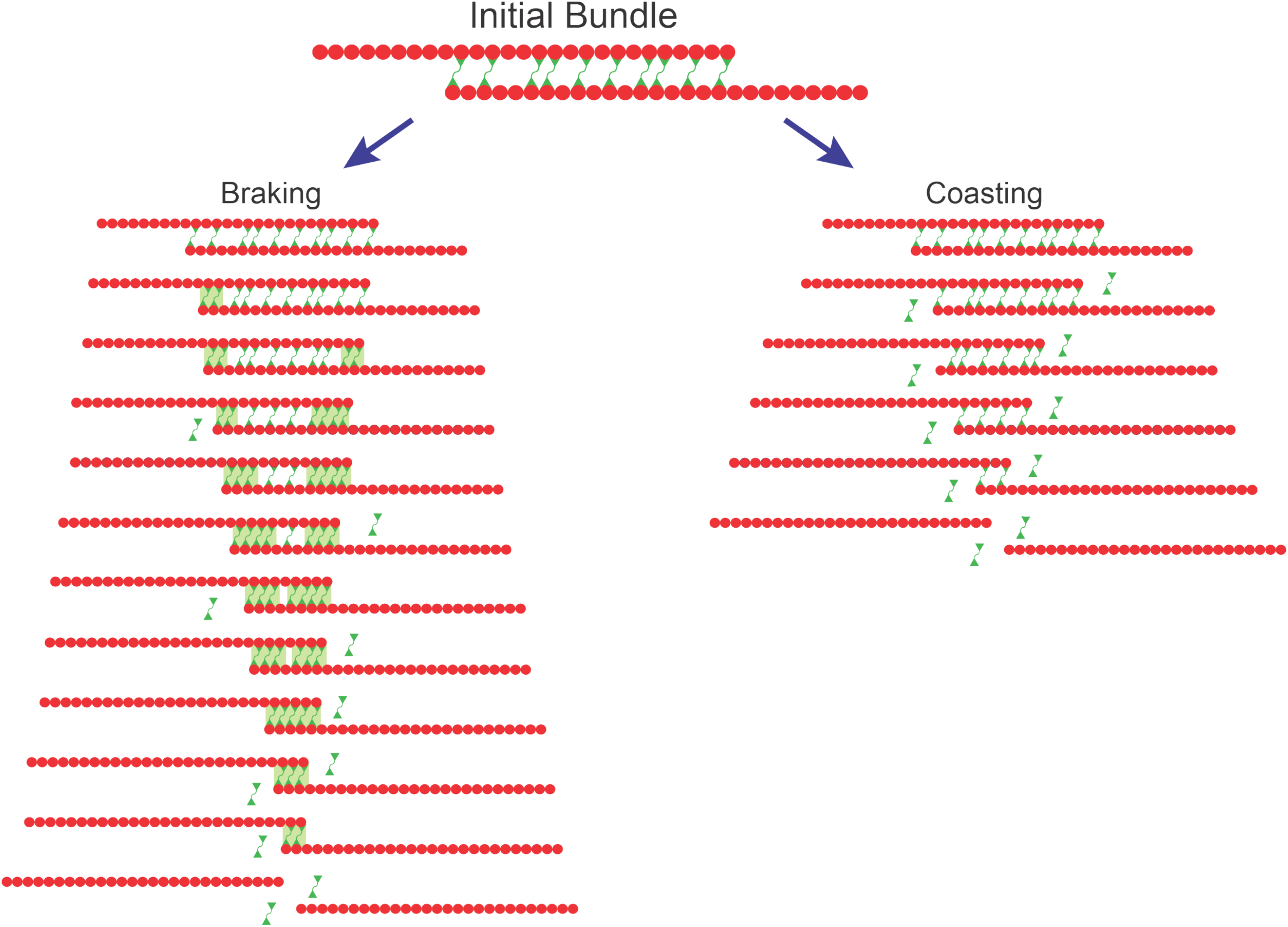
Model Describing Two Modes of Microtubule Sliding Determined by PRC1 Behavior. Schematized model depicting evolution of braking (left) and coasting (right) event bundles. Both bundles contain the same initial amount of PRC1 (green). Red circles represent 8 nm-long PRC1 binding sites on the microtubule lattice. Braking events slide apart more slowly than coasting events and retain more PRC1 over time. PRC1 forms clusters (green boxes) at the overlap edges that accumulate additional molecules as time increases. Coasting events have a shorter duration than braking events, losing PRC1 from the overlap more readily and lack PRC1 cluster formation at the edges.

Our results indicate that PRC1-mediated crosslinking of microtubule pairs always slows their gliding velocities relative to single filament rates, regardless of whether coasting or braking behavior emerges. Coasting provides resistance to motor-driven sliding via an ensemble of PRC1 crosslinking bridges that remain evenly distributed and likely highly diffusive within overlaps. We recently reported that mechanical separation of bundles at constant velocity and higher ionic strengths using an optical tweezers instrument generates viscous resistive forces that scale with pulling velocity (Gaska et al., 2020). These results are consistent with PRC1 adopting a coasting-like behavior, with uniform distribution of diffusive crosslinker particles throughout the overlap region. We hypothesize that this coasting mode would allow for the integration of diverse motor activity by resisting faster motor proteins more than slower proteins. We also observed that when sliding is paused and then resumed, the resulting resistance is greater, possibly due to clustering of PRC1 particles (Gaska et al., 2020). We speculate that these observations are consistent with a transition into the braking mode observed in this study. It is likely that coasting and braking behaviors are both important to the regulation of spindle dynamics, and the interplay of these two modes is likely spatiotemporally regulated. The proper distribution and arrangement of PRC1 molecules may be essential in determining the capacity of a bundle to resist motor-driven sliding.

How might such a distinct braking state emerge, and what might be the molecular arrangement of PRC1 molecules near tips? We hypothesize that in braking events, a cooperative structure of PRC1 molecules may spontaneously form near tips that is much less diffuse, stiffer, and less likely to detach from the microtubule surface. Previous cryo-EM work showed that PRC1 adopts different conformations based on its density within microtubule overlaps. Tightly packed PRC1 molecules form a highly-ordered structure with regular 8nm spacing between molecules, while sparser PRC1 molecules remain more diffusive (Subramanian et al., 2010). Therefore, it is possible that at high PRC1 density, cooperative regularly-packed structures form with properties such as increased structural rigidity that strongly inhibit microtubule sliding. It is also possible that, at high density, rather than forming a cooperative structure, PRC1 crowding leads to a loss of potential microtubule binding sites which results in a reduced effective diffusion constant and increased resistance to sliding. Similar types of behavior have been observed with the yeast PRC1-homolog, Ase1. Ase1’s adaptive braking mechanism relies on the compaction of Ase1 in the overlap region, where the microtubule ends act as barriers to diffusion and resistance to filament separation takes the form of an entropic expansion force (Braun et al., 2017; Lansky et al., 2015). A third possibility is that PRC1 molecules form higher-order complexes due to increased local density and a decrease in available binding sites, but are not packing into a strictly regular array. Indeed, single diffusive Ase1 particles have been shown to interact to form static multimeric structures upon reaching a critical density on microtubules, and these multimers localized to microtubule overlaps (Kapitein et al., 2008). Therefore, we hypothesize that the clusters observed in braking events are the result of microtubule ends acting as barriers to PRC1 diffusion or dissociating, leading to dense packing PRC1 molecules at the overlap edges by one of the three mechanisms mentioned above. These high-density regions are then capable of significantly inhibiting microtubule sliding. In contrast, in coasting events PRC1 is likely more evenly distributed throughout the overlap region, allowing it to remain more diffusive and leave the overlap more readily, thereby providing less resistance against motor-driven sliding.

What role might a braking mode play in regulating the assembly of the spindle midzone? Our recent results suggest that viscous resistance to microtubule sliding during coasting-like modes can scale with velocity (Gaska et al., 2020), but cannot explain how stable overlaps with minimal relative filament sliding can be established. Here, we demonstrate that the spontaneous sliding-induced formation of high-density clusters of PRC1 alone produce a new mechanical state which is significantly harder to disrupt when pulling forces are applied to microtubules. Establishing such high densities of PRC1 molecules within midzone overlaps in dividing cells would likely require the activity of motor proteins, and not simply PRC1 alone. Indeed, ensembles of proteins containing PRC1 complexed with its binding partner, the kinesin-4 motor protein Kif4A, have been previously reported to be able to both form length-dependent end-tags at filament tips and to modulate microtubule sliding behavior to generate stable overlaps (Subramanian et al., 2013; Wijeratne and Subramanian, 2018). Similar slowing of filament sliding has been observed with the kinesin-14 HSET and Ase1, where Ase1 has been observed to form high-density clusters within overlaps and generate an adaptive braking mechanism that resists kinesin-14-driven microtubule sliding (Braun et al., 2011, 2017). Our results indicate that PRC1 alone can generate multiple modes of resistance to filament sliding forces, but that the spatial distribution of molecules needed to engage in braking likely requires other factors within the dividing cell.

We propose that the differing local PRC1 environments in microtubule overlaps and the development and distribution of these high-density substructures contribute to the generation of a passive resistive force, allowing PRC1 to act as a true molecular brake. While coasting behavior would contribute to regulation of the rate of pole separation, the transition to braking behavior would provide enhanced resistance upon reaching critical density that would favor the formation of stable overlaps. Therefore, both coasting and braking behavior could be employed within dividing cells and may be temporally regulated to modulate microtubule sliding throughout mitosis.

## Materials and Methods

### Protein expression and purification

#### PRC1

Full-length human PRC1 Isoform 2 was cloned into a pET-DUET plasmid containing an N-terminal histidine tag followed by a Tobacco Etch Virus (TEV) cleavage site. An eGFP sequence was inserted between the TEV cleavage site and the N-terminus of PRC1 with a ‘AAA’ linker sequence just after the eGFP. Proteins were expressed via BL21(DE3) Rosetta *Escherichia coli* cells (Novagen). Full-length PRC1 was expressed for 3.5 hours at 18°C after induction with 0.5 mM IPTG. Cells were lysed via sonication in lysis buffer (1 mg/mL lysozyme, 50 mM phosphate (pH 8.0), 10 mM imidazole, 1% Igepal and HALT protease inhibitor (Pierce), 2 mM TCEP Bond Breaker, 2 mM Benz-HCl, 1 mM PMSF). The lysate was clarified by ultracentrifugation and the supernatant was incubated with Ni-NTA for 1 hour at 4°C (G Biosciences). The resin was then washed with Wash Buffer (50 mM phosphate, (pH 8.0), 500 mM KCl, 10 mM imidazole, 0.1% tween, 0.5 mM TCEP, 1 mM PMSF) and the protein was then eluted using Elution Buffer (50 mM phosphate (pH 7.0), 250 mM imidazole, 150 mM KCl, 0.5 mM TCEP). Elute was pooled and concentrated to a volume of approximately 1 mL. Next, 1/30 w/w Pro TEV protease (PROMEGA), 1 mM DTT, and 50 uL of Pro TEV 20X buffer were added and the sample incubated in a 30°C water bath for 15 minutes before overnight dialysis at 4°C in Gel Filtration Buffer (1X BRB80 (pH 6.8), 150 mM KCl, 10 mM bME). Size exclusion chromatography (Superose-6 increase 10/300 column, GE Healthcare) was then performed in Gel Filtration Buffer with a Shimadzu High Performance Liquid Chromatograph. After collecting peak fractions and concentrating to ~0.5 mg/mL, sucrose was added to 35% w/v before flash freezing in liquid nitrogen.

#### Kinesin-1 K439

Plasmid encoding a truncated kinesin-1 construct (K439) with a C-terminally fused EB1 sequence to create stable dimers and a His_8_ tag (Woll et al., 2018) was generously donated by the Dr. Susan Gilbert lab. The protein expression protocol was identical to PRC1, except induction occurred overnight for 18 hours at 16°C. The pellet was resuspended in lysis buffer (10 mM NaPO_4_ (pH 7.2), 300 mM NaCl, 2 mM MgCl_2_, 0.1 mM EGTA, 10 mM PMSF, 1 mM DTT, 0.2 mM ATP, 1 mg/mL lysozyme, 30 mM imidazole, 3 uL benzonase (Novagen)) and sonicated. The lysate was clarified by ultracentrifugation and the supernatant incubated with Ni-NTA for 1 hour at 4°C (G Biosciences). The resin was then washed with Wash Buffer (20 mM NaPO_4_ (pH 7.2), 300 mM NaCl, 2 mM MgCl_2_, 0.1 mM EGTA, 0.02 mM ATP, 50 mM Imidazole) and the protein was then eluted using Elution Buffer (20 mM NaPO_4_ (pH 7.2), 300 mM NaCl, 2 mM MgCl_2_, 0.1 mM EGTA, 1 mM DTT, 0.02 mM ATP, 400 mM imidazole). Elute was pooled and concentrated to a volume of approximately 1 mL. The protein was then dialyzed overnight at 4°C in Dialysis Buffer (20 mM HEPES (pH 7.2), 300 mM NaCl, 0.1 mM EGTA, 0.1 mM EDTA, 5 mM MgAc, 50 mM KAc, 1 mM DTT). The next day the protein was dialyzed for 1 hour at 4°C in Buffer 1 (20 mM HEPES (pH 7.2), 200 mM NaCl, 0.1 mM EGTA, 0.1 mM EDTA, 5 mM MgAc, 50 mM KAc,1 mM DTT) followed by dialysis for 1 hour at 4°C in Buffer 2 (20 mM HEPES (pH 7.2), 150 mM NaCl, 0.1 mM EGTA, 0.1 mM EDTA, 5 mM MgAc, 50 mM KAc,1 mM DTT) Further purification was performed using size exclusion chromatography (Superose-6 increase, GE Healthcare) using Shimadzu High Performance Liquid Chromatograph. Peak fractions were pooled and dialyzed for 3 hours at 4°C in Buffer 3 (20 mM HEPES (pH 7.2), 100 mM NaCl, 0.1 mM EGTA, 0.1 mM EDTA, 5 mM MgAc, 50 mM KAc,1 mM DTT, 5% Sucrose). Protein was flash frozen in liquid nitrogen.

### Microtubule preparation

Microtubule tubulin reagents were purchased from Cytoskeleton, Inc. Microtubules for surface-immobilization were generated via mixture of rhodamine tubulin (TL590M) and unmodified tubulin at a ratio of 1:20 along with 1 mM GMPCPP. Microtubules were polymerized at 37°C for 20 minutes before clarification and stabilization in 20 μM Taxol following published protocols (Shimamoto et al., 2015).

### Flow Chamber Construction

The flow chamber design and assay preparation were modified from a previously described protocol (Shimamoto et al., 2015). Anti-parallel microtubule bundles were constructed using passivated glass coverslips coated with SVA-PEG at a ratio of 50 PEG:1 biotin-PEG. All reagents were prepared with 1X BRB80 buffer. PRC1 (2-8 nM) was mixed with microtubules and allowed to incubate while the chamber was being constructed. Following each reagent flow-in and incubation, a flush with ~2 chamber volumes of 1X BRB80 was performed. Reagents were introduced stepwise with the following order and incubation times: (1) 0.5 mg/mL neutravidin for 2 minutes; (2) 0.5 mg/mL alpha casein 5 minutes; (3) 0.01 mg/mL Anti-Histidine Antibody for 5 minutes; (4) 0.25 μg/mL K439 for 5 minutes; (6) PRC1-microtubule bundles were added to the final Reaction Buffer (0.5 mg/mL alpha casein, 20 μM Taxol, 1.25 mM MgCl_2_, 0.125 mM EGTA, Oxygen Scavenging System (4.5 mg/mL glucose, 350 U/mL glucose oxidase, 34 U/mL catalase, 1mM DTT), 10-500 μM ATP. The chamber was then sealed with clear nail polish prior to experiments.

### Image Acquisition

Microtubule bundles were imaged using two-channel TIRF microscopy using the following laser lines and exposure times: GFP-tagged PRC1 molecules were excited using a 488 nm laser (60% power, 200 ms exposure); and rhodamine microtubules were excited using a 561 nm laser (30% power, 100 ms exposure). Images were acquired using a Photometric Prime 95B camera controlled with Nikon NIS Elements software at overall acquisition rates of one frame per ~4 seconds. The stage was moved laterally in the X direction to collect data over two fields of view (total size: 244.2 x 132.0 μm) to increase the total observable field of view. Prior to analysis, images were visually screened to ensure that there were only two microtubules per bundle and there were no additional interactions with microtubules from other bundles.

### Image Analysis

Analysis of fluorescent data and generation of intensity linescan data sets were performed using a combination of FIJI (ImageJ) tools and custom-written Python software. For each pair of microtubules, a linescan with selection width of 7 pixels was drawn along the entire path of bundle separation in FIJI. The fluorescence intensity data along this line was saved for each frame, resulting in a 2D array of intensity values at all recorded time points. Using a custom-written Python script, the slope of rhodamine intensity changes in the last frame were calculated. A peak detection algorithm was used to find the largest changes in slope, which denoted the ends of the separated microtubules. Microtubule lengths were calculated from these points, which allowed for back calculation of the overlap length and identification of the overlap region in preceding frames. Calculations could then be performed on the GFP signal in these regions, including a summation over GFP intensity values in this region to determine integrated signal intensity and the variance of the GFP signal.

The peak detection algorithm was also applied to the GFP channel within the overlap regions. All identified peaks were fitted to a Gaussian distribution, which allowed for the calculation of the full width at half-maximum (FWHM) for each peak. An empirically-determined maximum peak width value was used to filter the results and eliminate false peaks. From this data, the GFP signal peak position, height, width in microns, integrated intensity, and GFP peak density were extracted.

## Acknowledgments

We would like to thank Dr. Susan Gilbert (RPI) for the gift of the kinesin-1 K439 plasmid and Dr. Blanca Barquera (RPI) for assistance with construct design. We also wish to thank Drs. Susan Gilbert, Alexey Khodjakov, Haixin Sui, Rebecca Fisher (Wadsworth Center) and Marvin Bentley (RPI) for providing advice and suggestions for experiments and data analysis methods. We also wish to thank members of the Forth lab for helpful discussions and critical reading of the manuscript. This work was supported by startup funds to S.F. provided by the School of Science at Rensselaer Polytechnic Institute.

## Author contributions

A.A. and S.F. conceived of experimental design. A.A. performed data acquisition and analysis. A.A. and I.G. performed expression and purification of proteins used in the study. A.A. and S.F. wrote and edited manuscript with assistance from I.G.

## Supplemental Figures S1-S5

**Figure S1:**
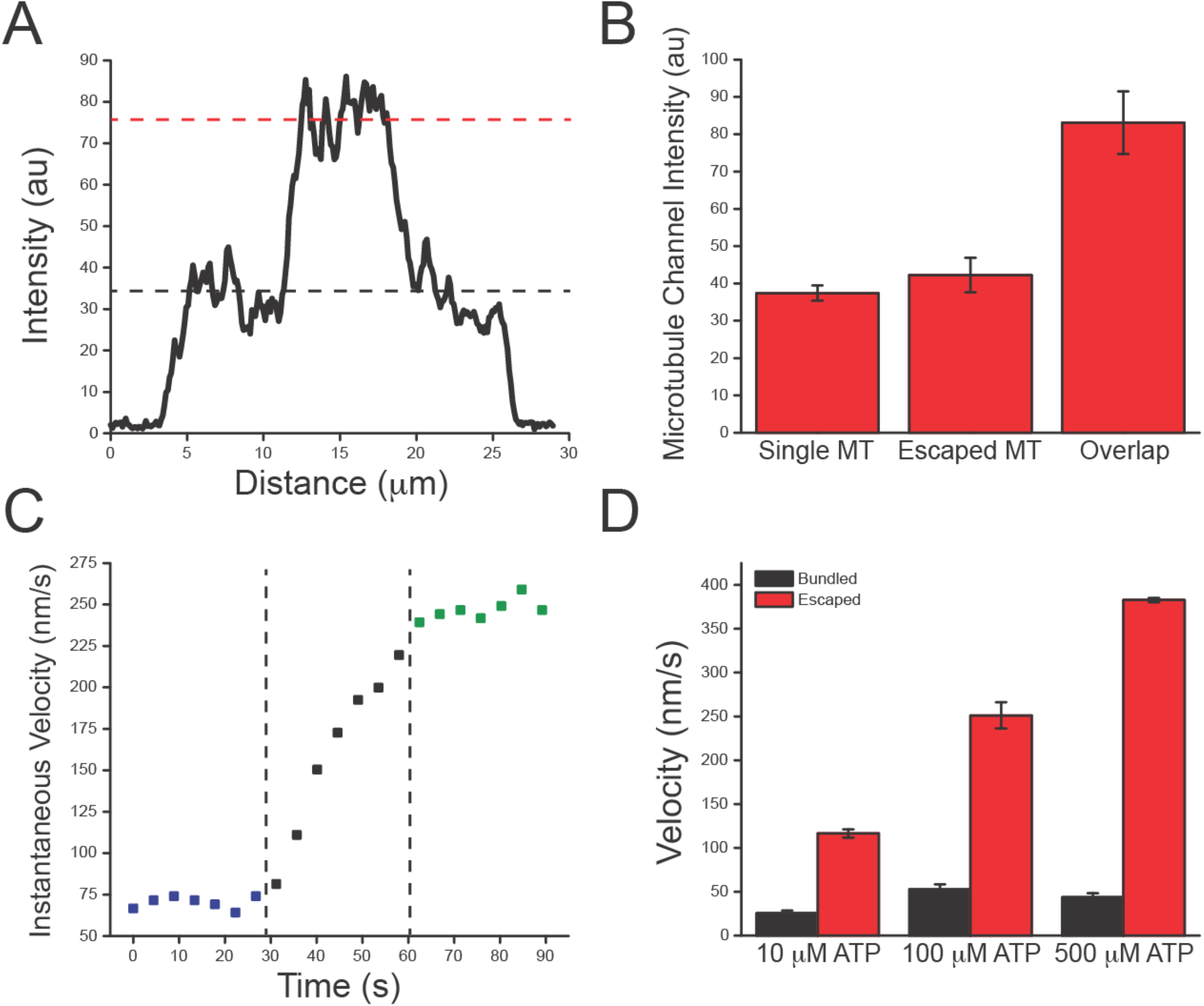
PRC1 Slows microtubule Sliding Over a Range of Kinesin Stepping Rates. (A) Sample microtubule fluorescence intensity linescan of a bundle. Dashed red line depicts average intensity of two overlapping microtubules. Black line denotes average intensity of single microtubules. (B) Quantification of average microtubule intensities of single, escaped, and overlapping microtubules. Single microtubules were never part of a bundle. N = 25 (Single), 13 (Escaped), 13 (Overlap). (C) A representative example of calculated microtubule velocities as the bundle slides apart. Bundled microtubules (blue) have lower velocities than escaped microtubules (green), with an acceleration phase (black). (D) Bundled (black) and escaped (red) microtubule velocities at various ATP concentrations. Escaped microtubule velocities are consistently higher than bundled velocities. N = 48, 33 (10 μM ATP), 42, 46 (100 μM ATP), 26, 21 (500 μM ATP) for braking and coasting, respectively. All error bars are SEM.

**Figure S2:**
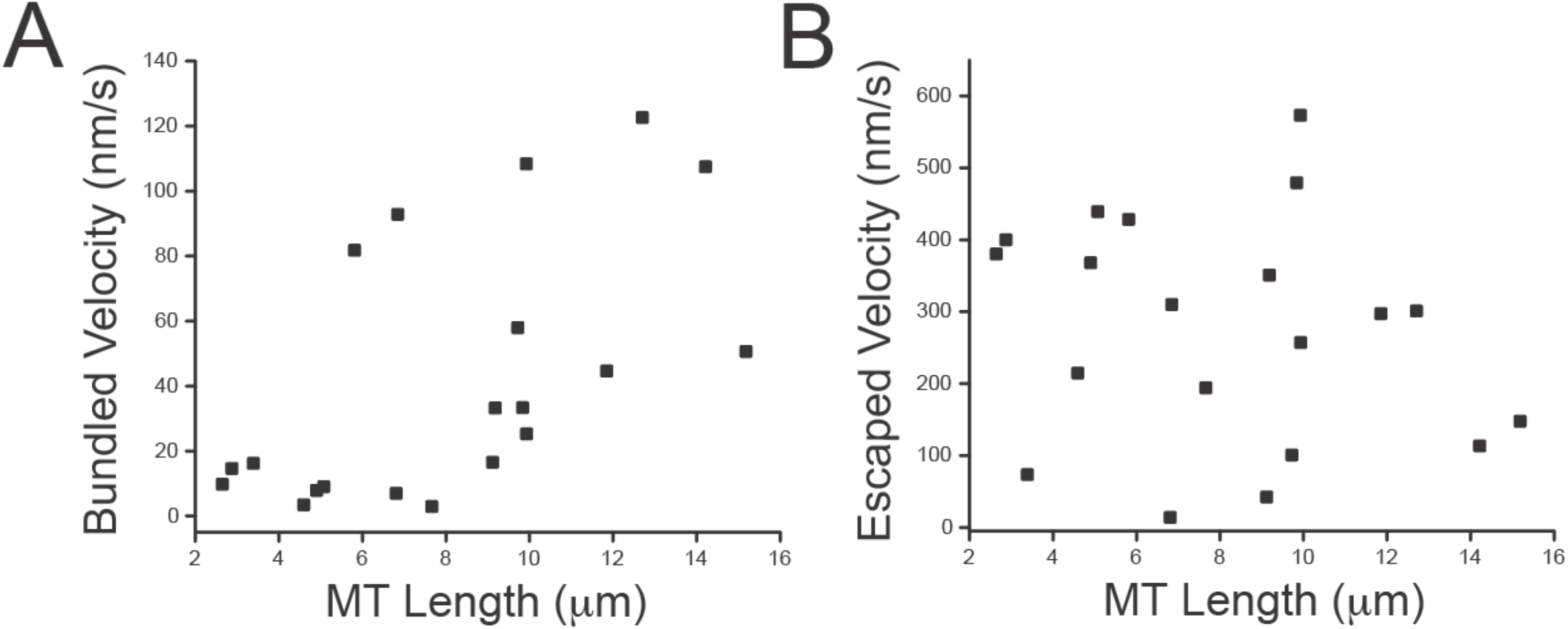
Longer Microtubules Have Slightly Higher Bundled Velocities. (A) Bundled microtubule velocities exhibit a slight positive correlation with increasing microtubule length. (B) Escaped microtubule velocities exhibit no dependence on microtubule length. N = 20 for both (A) and (B).

**Figure S3:**
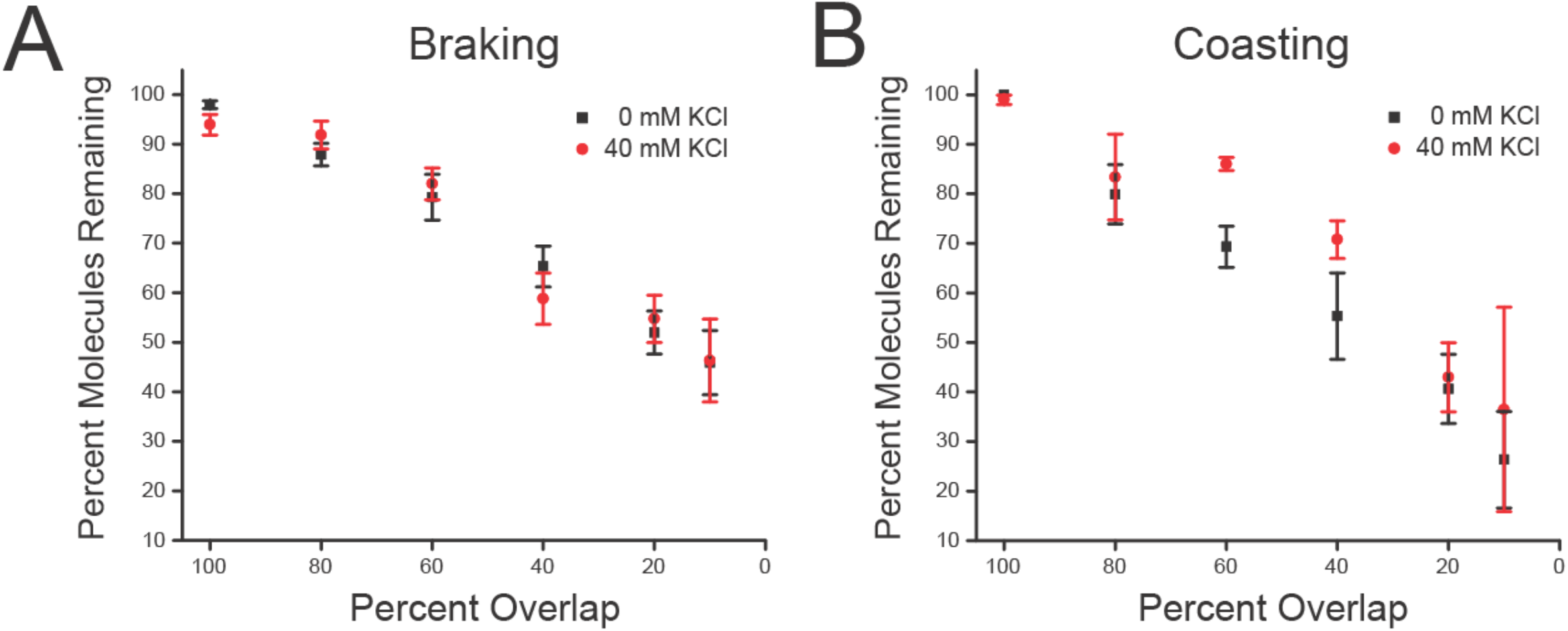
Increasing Ionic Strength Does Not Impact PRC1 Retention in Overlaps. (A) Percentage of PRC1 molecules remaining as a function of decreasing overlap length for braking events with 0 (black) or 40 (red) mM additional KCl in the final buffer solution at 100 μM ATP. Number of bundles averaged for each point (40 mM KCl): 12 (100%), 7 (80%), 9 (60%), 9 (40%), 10 (20%), 7 (10%). Number of bundles averaged for each point for 0 mM KCl are the same as in 3C. (B) Percentage of PRC1 molecules remaining as a function of decreasing overlap length for coasting events with 0 (black) or 40 (red) mM additional KCl. Number of bundles averaged for each point (40 mM KCl): 8 (100%), 4 (80%), 4 (60%), 2 (40%), 6 (20%), 2 (10%). Number of bundles averaged for each point (0 mM KCl) are the same as in 3C. All error bars are SEM.

**Figure S4:**
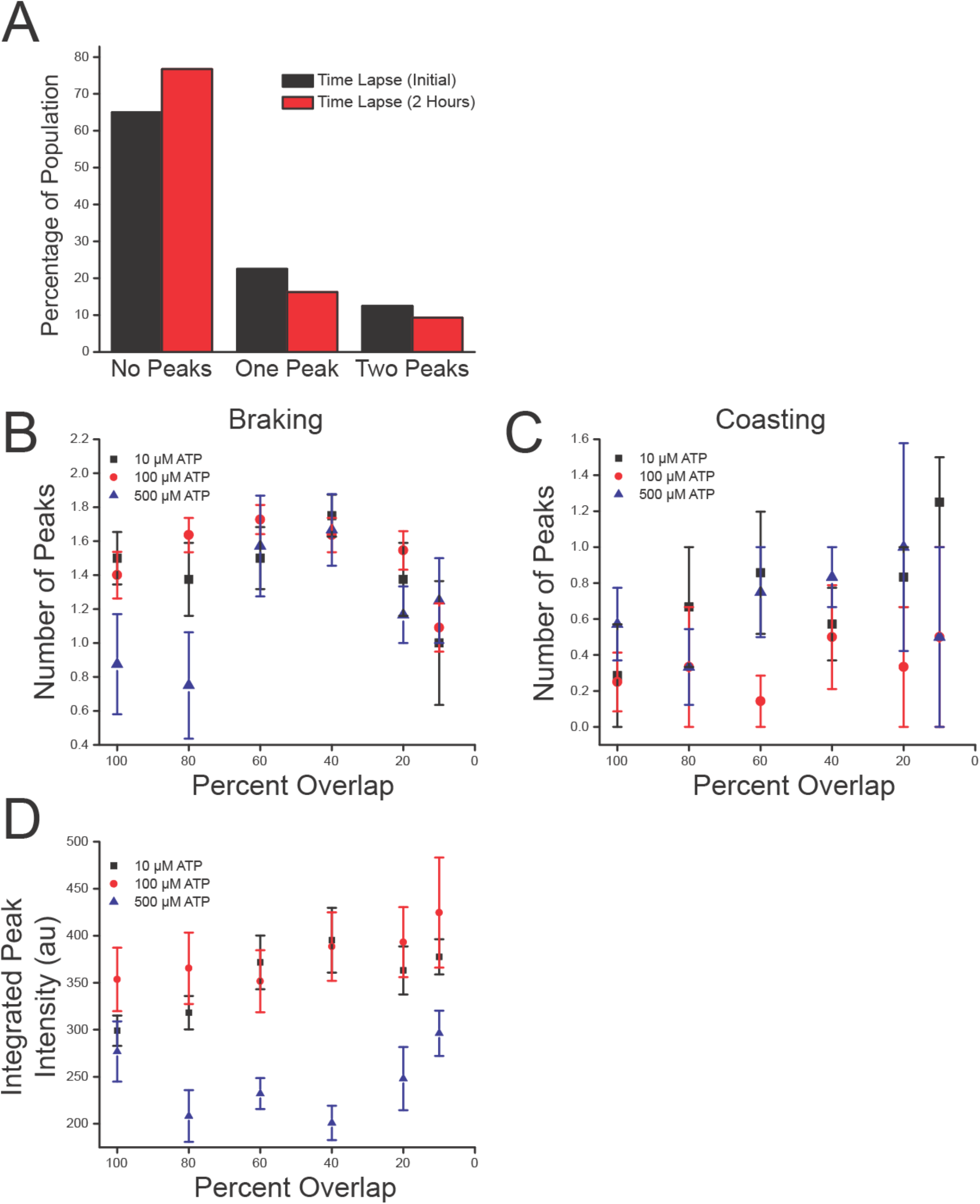
PRC1 Forms Clusters Across a Range of Kinesin Stepping Rates. (A) Number of peaks present in a population of static PRC1-microtubule bundles (bound to surface-kinesins but without the addition of ATP) after a 20 minute incubation (black, “initial”) and two hours later (red). At both time points the majority of bundles contain no peaks, and this percentage increases after two hours. N = 26, 33 (No Peaks), 9, 7 (One Peak), 5,4 (Two Peaks) for Initial and 2 Hour time points, respectively. (B) Number of GFP-PRC1 peaks as a function of decreasing overlap length for braking event overlaps at 10 (black), 100 (red), and 500 (blue) μM ATP. Number of individual peaks averaged for each point: 12, 14, 14 (100%), 12, 18, 12 (80%), 9, 19, 12 (60%), 14, 18, 12 (40%), 12, 17, 10 (20%), 5, 12, 7 (10%) for 10, 100, and 500 μM ATP, respectively. 100 μM data replotted from (5I). (C) Number of GFP-PRC1 peaks as a function of decreasing overlap length for coasting event overlaps at 10 (black), 100 (red), and 500 (blue) μM ATP. Number of individual peaks averaged for each point: 7, 8, 9 (100%), 6, 3, 9 (80%), 7, 7, 8 (60%), 7, 4, 7 (40%), 6, 3, 7 (20%), 4, 2, 4 (10%) for 10, 100, and 500 μM ATP, respectively. 100 μM data replotted from (5I). (D) Integrated peak intensities as a function of decreasing overlap length for braking event overlaps at 10 (black), 100 (red), and 500 (blue) μM ATP. Number of individual peaks averaged for each point are the same as in (B), except for 10 μM at 80% (N = 11), 20% (N = 11) and 10% (N = 4).100 μM data replotted from (5J). All error bars are SEM.

**Figure S5:**
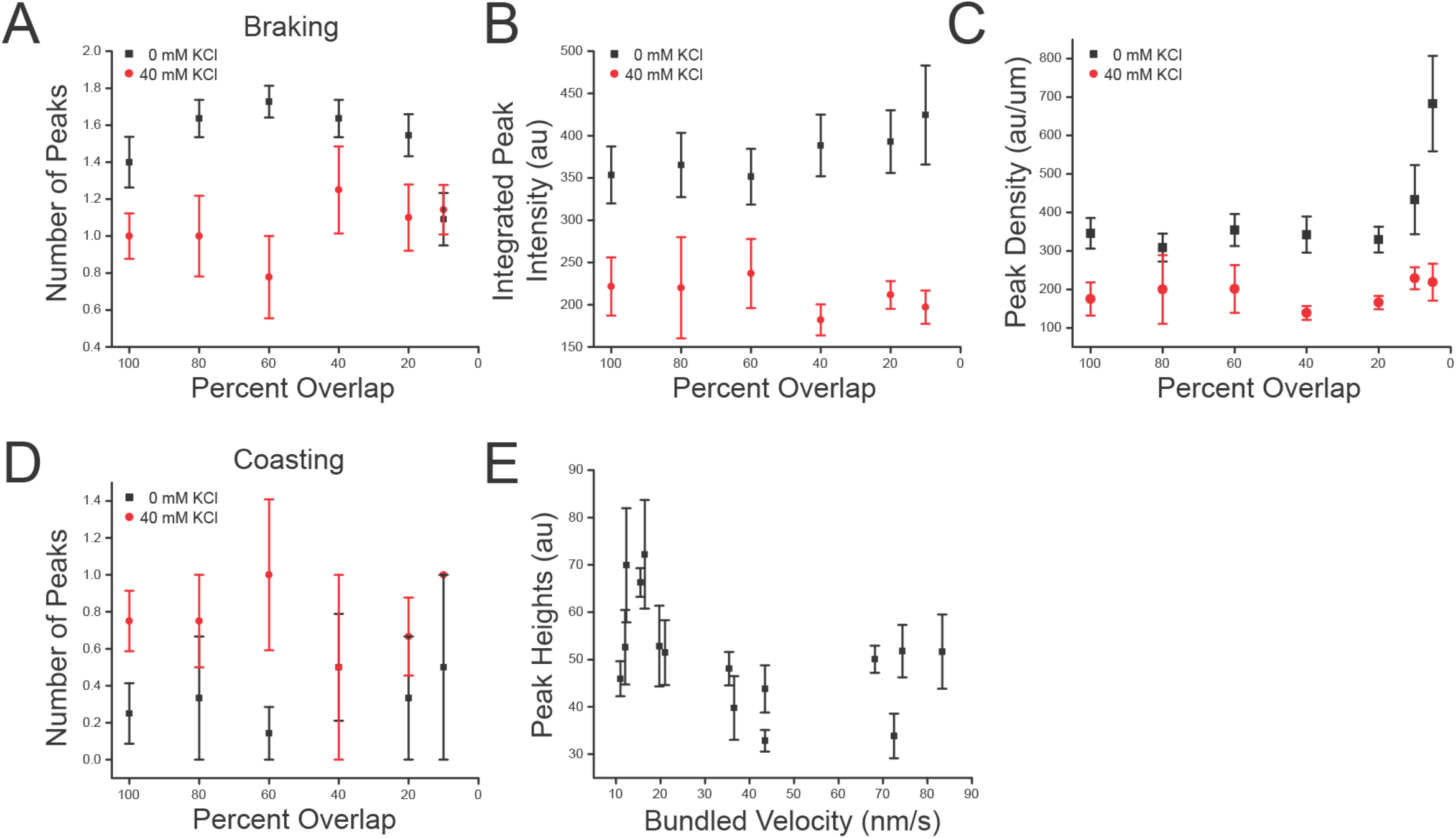
Increasing Ionic Strength Impairs PRC1 Cluster Formation. (A) Number of peaks in GFP-PRC1 intensity as a function of decreasing overlap length for braking events with 0 (black) and 40 (red) mM additional KCl in the final buffer solution at 100 μM ATP. Number of individual peaks averaged for each point (40 mM KCl): 12 (100%), 7 (80%), 7 (60%), 10 (40%), 11 (20%), 8 (10%). Number of individual peaks averaged for 0 mM KCl are the same as in 5I. 0 mM data replotted from Fig. 5I. (B) Integrated peak intensity as a function of decreasing overlap length for braking event peaks with 0 (black) and 40 (red) mM additional KCl at 100 μM ATP. Number of individual peaks averaged for each point are the same as in (A). 0 mM data replotted from Fig. 5J. Peak density as a function of decreasing overlap length for braking event peaks with 0 (black) and 40 (red) mM additional KCl at 100 μM ATP. Number of individual peaks averaged for 40 mM KCl are the same as in (B). N for 5% overlap length = 3. Number of individual peaks averaged for 0 mM KCl are the same as in 5K. 0 mM KCl data replotted from 5K. All error bars are SEM. (D) Number of peaks in GFP-PRC1 intensity as a function of decreasing overlap length for coasting events with 0 (black) and 40 (red) mM additional KCl at 100 μM ATP. Number of individual peaks averaged for each point (40 mM KCl): 8 (100%), 4 (80%), 4 (60%), 2 (40%), 6 (20%), 3 (10%). Number of individual peaks averaged for 0 mM KCl are the same as in 5I. 0 mM data replotted from 5I. (E) Peak heights show a slight negative correlation with increasing bundled velocity in braking events. Each point represents one bundle (N = 15). Peak heights for a given bundle are averaged (N = 2). All error bars are SEM.

## References

Bieling P, Kandels-Lewis S, Telley IA, Van Dijk J, Janke C, Surrey T. 2008. CLIP-170 tracks growing microtubule ends by dynamically recognizing composite EB1/tubulinbinding sites. J Cell Biol 183:1223–1233. doi: 10.1083/jcb.200809190

Bieling P, Telley IA, Surrey T. 2010. A minimal midzone protein module controls formation and length of antiparallel microtubule overlaps. Cell 142:420–432. doi:10.1016/j.cell.2010.06.033

Braun M, Lansky Z, Fink G, Ruhnow F, Diez S, Janson ME. 2011. Adaptive braking by Ase1 prevents overlapping microtubules from sliding completely apart. Nat Cell Biol 13:1259–1264. doi:10.1038/ncb2323

Braun M, Lansky Z, Szuba A, Schwarz FW, Mitra A, Gao M, Lüdecke A, Ten Wolde PR, Diez S. 2017. Changes in microtubule overlap length regulate kinesin-14-driven microtubule sliding. Nat Chem Biol 13:1245–1252. doi:10.1038/nchembio.2495

Duellberg C, Trokter M, Jha R, Sen I, Steinmetz MO, Surrey T. 2014. Reconstitution of a hierarchical +TIP interaction network controlling microtubule end tracking of dynein. Nat Cell Biol 16:804–811. doi:10.1038/ncb2999

Fallesen T, Roostalu J, Duellberg C, Pruessner G, Surrey T. 2017. Ensembles of Bidirectional Kinesin Cin8 Produce Additive Forces in Both Directions of Movement. Biophys J 113:2055–2067. doi:10.1016/j.bpj.2017.09.006

Forth S, Hsia KC, Shimamoto Y, Kapoor TM. 2014. Asymmetric friction of nonmotor MAPs can lead to their directional motion in active microtubule networks. Cell 157:420–432. doi:10.1016/j.cell.2014.02.018

Forth S, Kapoor TM. 2017. The mechanics of microtubule networks in cell division. J Cell Biol. doi:10.1083/jcb.201612064

Gaska I, Armstrong M, Alfieri A, Forth S. 2020. The Mitotic Crosslinking Protein PRC1 Acts as a Mechanical Dashpot to Resist Microtubule Sliding. Dev Cell 54:1–14. doi:10.2139/ssrn.3481313

Janson ME, Loughlin R, Loïodice I, Fu C, Brunner D, Nédélec FJ, Tran PT. 2007. Crosslinkers and Motors Organize Dynamic Microtubules to Form Stable Bipolar Arrays in Fission Yeast. Cell 128:357–368. doi:10.1016/j.cell.2006.12.030

Jonsson E, Yamada M, Vale RD, Goshima G. 2015. Clustering of a kinesin-14 motor enables processive retrograde microtubule-based transport in plants. Nat Plants 1:1–7. doi:10.1038/nplants.2015.87

Kajtez J, Solomatina A, Novak M, Polak B, Vukušić K, Rüdiger J, Cojoc G, Milas A, Šumanovac Šestak I, Risteski P, Tavano F, Klemm AH, Roscioli E, Welburn J, Cimini D, Glunčić M, Pavin N, Tolić IM. 2016. Overlap microtubules link sister k-fibres and balance the forces on bi-oriented kinetochores. Nat Commun 7:11. doi:10.1038/ncomms10298

Kapitein LC, Janson ME, van den Wildenberg SMJL, Hoogenraad CC, Schmidt CF, Peterman EJG. 2008. Microtubule-Driven Multimerization Recruits ase1p onto Overlapping Microtubules. Curr Biol 18:1713–1717. doi:10.1016/j.cub.2008.09.046

Kapoor T. 2017. Metaphase Spindle Assembly. Biology (Basel) 6:8. doi:10.3390/biology6010008

Lansky Z, Braun M, Lüdecke A, Schlierf M, Ten Wolde PR, Janson ME, Diez S. 2015. Diffusible crosslinkers generate directed forces in microtubule networks. Cell 160:1159–1168. doi:10.1016/j.cell.2015.01.051

Maurer SP, Fourniol FJ, Bohner G, Moores CA, Surrey T. 2012. EBs recognize a nucleotide-dependent structural cap at growing microtubule ends. Cell 149:371–382. doi:10.1016/j.cell.2012.02.049

McIntosh J, Hays T. 2016. A Brief History of Research on Mitotic Mechanisms. Biology (Basel) 5:55. doi:10.3390/biology5040055

Mollinari C, Kleman JP, Jiang W, Schoehn G, Hunter T, Margolis RL. 2002. PRC1 is a microtubule binding and bundling protein essential to maintain the mitotic spindle midzone. J Cell Biol 157:1175–1186. doi:10.1083/jcb.200111052

Mollinari C, Kleman JP, Saoudi Y, Jablonski SA, Perard J, Yen TJ, Margolis RL. 2005. Ablation of PRC1 by small interfering RNA demonstrates that cytokinetic abscission requires a central spindle bundle in mammalian cells, whereas completion of furrowing does not. Mol Biol Cell 16:1043–1055. doi:10.1091/mbc.E04-04-0346

Pamula MC, Carlini L, Forth S, Verma P, Suresh S, Legant WR, Khodjakov A, Betzig E, Kapoor TM. 2019. High-resolution imaging reveals how the spindle midzone impacts chromosome movement. J Cell Biol 218:2529–2544. doi:10.1083/jcb.201904169

Pecreaux J, Röper JC, Kruse K, Jülicher F, Hyman AA, Grill SW, Howard J. 2006. Spindle Oscillations during Asymmetric Cell Division Require a Threshold Number of Active Cortical Force Generators. Curr Biol 16:2111–2122. doi:10.1016/j.cub.2006.09.030

Polak B, Risteski P, Lesjak S, Tolić IM. 2017. PRC 1-labeled microtubule bundles and kinetochore pairs show one-to-one association in metaphase. EMBO Rep 18:217–230. doi:10.15252/embr.201642650

Rincon SA, Lamson A, Blackwell R, Syrovatkina V, Fraisier V, Paoletti A, Betterton MD, Tran PT. 2017. Kinesin-5-independent mitotic spindle assembly requires the antiparallel microtubule crosslinker Ase1 in fission yeast. Nat Commun 8. doi:10.1038/ncomms15286

Roostalu J, Hentrich C, Bieling P, Telley IA, Schiebel E, Surrey T. 2011. Directional switching of the kinesin Cin8 through motor coupling. Science (80-) 332:94–99. doi:10.1126/science.1199945

Shapira O, Goldstein A, Al-Bassam J, Gheber L. 2017. A potential physiological role for bi-directional motility and motor clustering of mitotic kinesin-5 Cin8 in yeast mitosis. J Cell Sci 130:725–734. doi:10.1242/jcs.195040

Shimamoto Y, Forth S, Kapoor TM. 2015. Measuring Pushing and Braking Forces Generated by Ensembles of Kinesin-5 Crosslinking Two Microtubules. Dev Cell 34:669–681. doi:10.1016/j.devcel.2015.08.017

Subramanian R, Kapoor TM. 2012. Building Complexity: Insights into Self-Organized Assembly of Microtubule-Based Architectures. Dev Cell. doi:10.1016/j.devcel.2012.10.011

Subramanian R, Ti SC, Tan L, Darst SA, Kapoor TM. 2013. Marking and measuring single microtubules by PRC1 and kinesin-4. Cell 154:377–390. doi:10.1016/j.cell.2013.06.021

Subramanian R, Wilson-Kubalek EM, Arthur CP, Bick MJ, Campbell EA, Darst SA, Milligan RA, Kapoor TM. 2010. Insights into antiparallel microtubule crosslinking by PRC1, a conserved nonmotor microtubule binding protein. Cell 142:433–443. doi:10.1016/j.cell.2010.07.012

Tikhonenko I, Irizarry K, Khodjakov A, Koonce MP. 2016. Organization of microtubule assemblies in Dictyostelium syncytia depends on the microtubule crosslinker, Ase1. Cell Mol Life Sci 73:859–868. doi:10.1007/s00018-015-2026-8

Varga V, Helenius J, Tanaka K, Hyman AA, Tanaka TU, Howard J. 2006. Yeast kinesin-8 depolymerizes microtubules in a length-dependent manner. Nat Cell Biol 8:957–962. doi:10.1038/ncb1462

Varga V, Leduc C, Bormuth V, Diez S, Howard J. 2009. Kinesin-8 Motors Act Cooperatively to Mediate Length-Dependent Microtubule Depolymerization. Cell 138:1174–1183. doi:10.1016/j.cell.2009.07.032

Vukušić K, Buđa R, Bosilj A, Milas A, Pavin N, Tolić IM. 2017. Microtubule Sliding within the Bridging Fiber Pushes Kinetochore Fibers Apart to Segregate Chromosomes. Dev Cell 43:11–23.e6. doi:10.1016/j.devcel.2017.09.010

Wijeratne S, Subramanian R. 2018. Geometry of antiparallel microtubule bundles regulates relative sliding and stalling by PRC1 and kif4A. Elife 7. doi:10.7554/eLife.32595

Woll KA, Guzik-Lendrum S, Bensel BM, Bhanu N V., Dailey WP, Garcia BA, Gilbert SP, Eckenhoff RG. 2018. An allosteric propofol-binding site in kinesin disrupts kinesin-mediated processive movement on microtubules. J Biol Chem 293:11283–11295. doi:10.1074/jbc.RA118.002182

Yamashita A, Sato M, Fujita A, Yamamoto M, Toda T. 2005. The Roles of Fission Yeast Ase1 in Mitotic Cell Division, Meiotic Nuclear Oscillation, and Cytokinesis Checkpoint Signaling. Mol Biol Cell 16:1378–1395. doi:10.1091/mbc.e04-10-0859

Zhu C, Lau E, Schwarzenbacher R, Bossy-Wetzel E, Jiang W. 2006. Spatiotemporal control of spindle midzone formation by PRC1 in human cells. Proc Natl Acad Sci U S A 103:6196–201. doi:10.1073/pnas.0506926103

